# De novo design of protein-binding aptamers through deep reinforcement learning assembly of nucleic acid fragments

**DOI:** 10.1101/2025.06.01.657174

**Authors:** Gaoxing Guo, Liangwei Guo, Jiaqiang Qian, Xiaoming He, Xinzhou Qian, Lei Wang, Qiang Huang

## Abstract

Nucleic acid aptamers targeting proteins are becoming increasingly important in biopharmaceuticals and molecular diagnostics. Traditionally, aptamers are discovered through labor-intensive screening of nucleic acid libraries using the SELEX method. However, de novo design approaches without experimental screening remain a significant challenge. Here, we employed deep reinforcement learning to develop an artificial intelligence (AI) agent capable of de novo aptamer design, termed AiDTA (AI-driven Docking-Then-Assembling). First, nucleic acid fragments were docked to the target protein to identify target-binding fragments. Then, AiDTA automatically assembled these fragments into aptamers using the Monte Carlo tree search algorithm and a policy-value neural network to guide the agent in generating aptamers with secondary structures similar to the original constituent fragments. Experimental validation demonstrated that the AiDTA-designed DNA aptamers targeting disease-related proteins exhibited high binding affinities in the nanomolar range, achieving the de novo design of protein-binding aptamers for the first time. Our study establishes a new approach to obtaining protein-binding aptamers for potential applications in biopharmaceuticals and diagnostics.

## Introduction

Nucleic acid aptamers are single-stranded DNA or RNA molecules, typically ranging from 15 to 80 nucleotides (nt) in length, that can bind to specific targets with high affinity and specificity through their unique three-dimensional (3D) structures. These molecules can interact with small molecules^1,2^, metal ions^3^, proteins^4,5^, cells^6^, viruses^7^, and bacteria^8^. This versatility makes aptamers highly suitable for applications in biopharmaceuticals^9,10^, molecular diagnostics^11,12^, biosensors^13^, and drug delivery systems^14^. Despite such application potentials, aptamers are mainly discovered by screening large random sequence libraries of nucleic acids using the Systematic Evolution of Ligands by Exponential Enrichment (SELEX) technique *in vitro*, a method introduced more than three decades ago^15,16^. The SELEX screening is a resource-intensive and time-consuming process, involving iterative cycles of library construction, target binding, elution, separation, and amplification by PCR. Nucleic acid libraries used for screening typically contain 10^14^∼10^15^ random sequences of 20 to 80 nt^17^. Often, 8 to 15 rounds of SELEX screening are required, and each round takes several days to a week. In addition to being resource intensive, SELEX libraries sample only a very small fraction of the vast, possible sequence space, since the theoretical space for 80-nt sequences is about 10^48^. Therefore, the development of more efficient aptamer discovery or design methods is important to overcome the inherent limitations of the traditional SELEX method.

In recent years, artificial intelligence (AI) technologies have been increasingly applied to enhance the SELEX processes, with the aim of shortening its experimental cycles and efficiently discovering the high-affinity aptamers from the SELEX data. For example, Alipanahi et al. and Hassanzadeh et al. developed DeepBind^18^ and DeeperBind^19^, respectively, to predict the sequence specificities of DNA- and RNA-binding proteins, which is also able to detect the protein-binding sites of the SELEX aptamers. Pan et al. introduced iDeepS which can identify the potential sequence and structural motifs of aptamers^20^. Specifically for the SELEX screening, Iwano et al. developed a variational autoencoder RaptGen^21^ that learns from SELEX data to create a latent space in which sequences are clustered by motif structure, thereby generating aptamers that are not included in the SELEX data. Similarly, Wang et al. introduced AptaDiff^22^, which was also able to generate high-affinity aptamers not present in the SELEX data. Very recently, Wong et al. developed RhoDesign for generative design of RNA aptamers capable of fluorescing upon binding to small molecule TO1-biotin^23^. All these studies demonstrated the great potential of AI deep learning technologies to overcome the limitations of SELEX. However, these methods still require SELEX data to train the deep learning models, or depend on predetermined aptamer structures. So far, there are no general de novo aptamer design methods that do not rely on experimental screening or information.

In this respect, we have previously proposed the docking-then-assembling (DTA) method for de novo aptamer design and successfully applied it to develop DNA aptamers targeting the marine biotoxin okadaic acid (OA)^24^. The DTA method consists of three main steps: (1) identifying high-affinity binding sites on the target molecule for single nucleotides by molecular docking; (2) assembling the unlinked single nucleotides (hereafter defined as nucleic acid fragments) at these high-affinity binding sites into binding units; and (3) linking these binding units with stabilizing fragments to form the final full-length aptamers. Our DTA method demonstrated that aptamers could be designed for a specific target without using SELEX screening. However, this method relied heavily on manual assembly of the unconnected fragments into the full-length aptamers, and thus could only explore a very limited sequence space. As a result, only a very small number of candidate aptamers were designed, and in the design process the secondary structures that are critical for aptamer functionality^25^ were not sufficiently taken into account.

The successful applications of deep learning techniques in protein design inspired us to explore the integration of AI technology into the DTA method, in order to overcome its limitations of manual assembly. In the DTA method, an aptamer is designed by iteratively selecting fragments from a pool containing target-binding fragments and then linking them to form a continuous sequence of certain length. Fragment selection and the determination of their linkage positions in the assembling sequence can be considered as a decision-making action; a series of assembling actions (i.e., fragment selection and linkage positioning) will ultimately generate full-length aptamers. The objective of these assembling actions is to produce an aptamer whose target-binding modes closely resemble those of its constituent fragments. This fragment assembly process is inherently well-suited for reinforcement learning (RL), a framework in which a trainable AI agent interacts with an environment to learn an optimal decision-making policy^26^. In RL, the AI agent observes the current state of the environment, takes an action, and transitions to a new state while receiving a reward based on the outcome of the action; through iterative optimization of its action policies, the agent aims to maximize long-term cumulative rewards^27^. As illustrated in Fig. 1a, for the DTA approach, fragment selection and linkage positioning can be conceptualized as the actions of the AI agent, while the target molecule serves as the environment. The current state of the environment corresponds to the partially assembled aptamer sequence, and the reward is determined by the similarity between secondary structures of the assembled aptamer and those of its constituent fragments.

Deep reinforcement learning (DeepRL) has achieved significant success in tasks that require continuous decision-making and interaction with the environment^28^. Since 2016, DeepMind has developed a series of AI agents for playing Go games, including AlphaGo^29^, AlphaGo Zero^30^, and AlphaZero^31^; and, trained by DeepRL, these AI agents defeated the world champion Go players, demonstrating that the DeepRL methods can solve very challenging tasks. Recently, DeepSeek-R1 also demonstrated that RL can significantly enhance and optimize the logical reasoning and problem-solving capabilities of large language models^32^. As a result, DeepRL techniques have begun to find their applications in the design of biological macromolecules. In 2023, David Baker’s group introduced a “top-down design” RL method that successfully designed protein nanocages with specific spatial frameworks and stacking geometries^33^. This method used the Monte Carlo Tree Search (MCTS) algorithm to sample the action space of assembling the protein nanocage subunits and optimized the design strategies to generate the protein nanocages that met spatial geometric constraints. Almost at the same time, Yang et al. developed a protein design AI agent (EvoPlay) based on the DeepRL framework of AlphaZero^34^. EvoPlay treated the amino acid mutations in a protein sequence of given length as the selection actions of 20 amino acids, and then used DeepRL to optimize the mutation selection strategies. These two studies demonstrated that DeepRL is particularly suitable for the design of biological macromolecules with sequence and structural constraints.

**Fig. 1:**
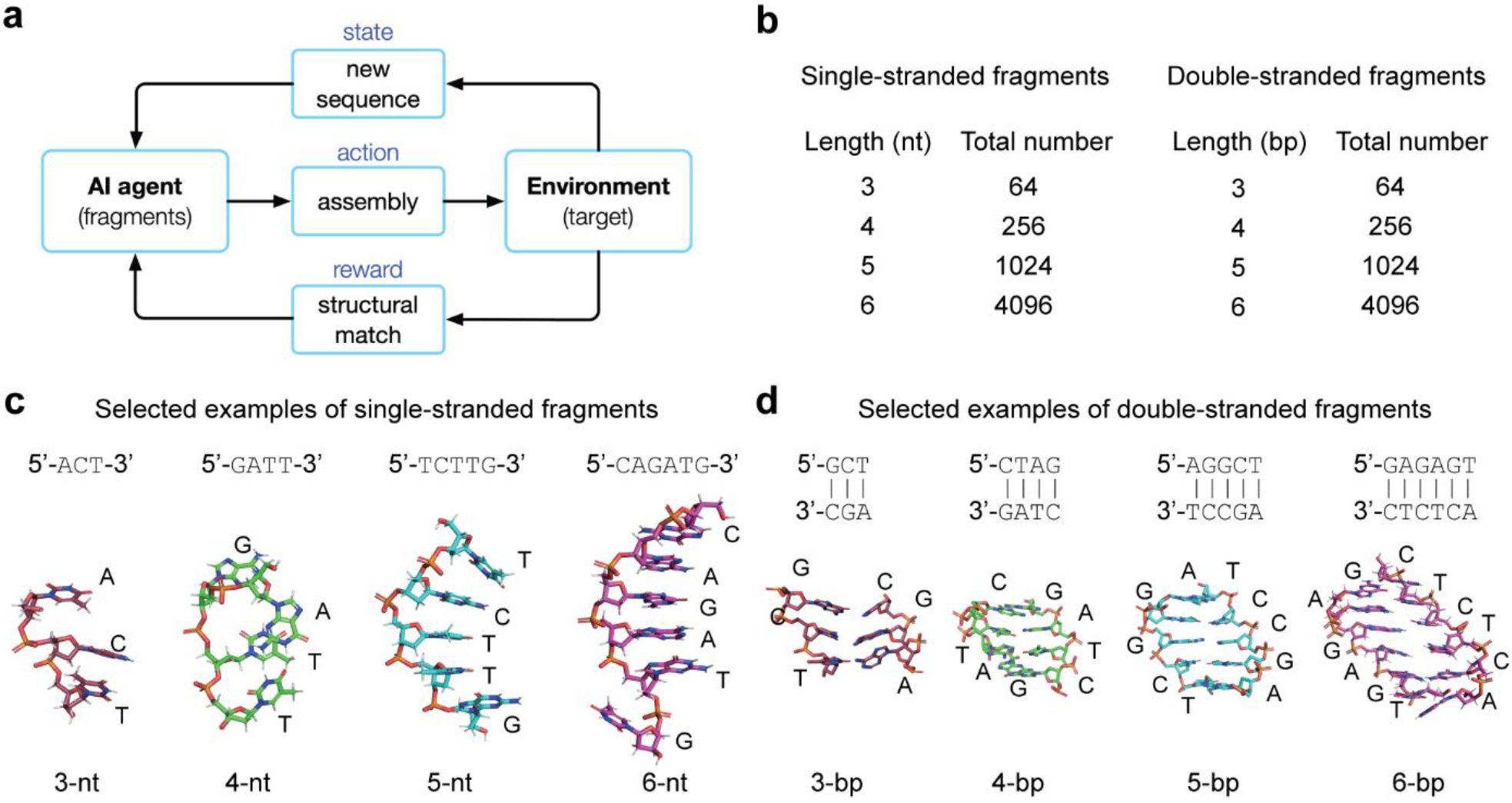
RL-driven fragment assembly and library of short nucleic acid fragments for molecular docking against protein targets. **a**, Schematic diagram of the RL-driven fragment assembly process. **b**, Total numbers of single-stranded and double-stranded fragments categorized by lengths. **c**, Representative 3D structures of the single-stranded fragments. **d**, Representative 3D structures of the double-stranded fragments.

In this study, we integrate DeepRL with the DTA approach to develop an AI agent capable of de novo design of protein-binding aptamers. First, we constructed a library of short single-stranded and double-stranded nucleic acid fragments, and then docked these fragments to the target protein to obtain a pool of high-affinity target-binding fragments. Next, we built an AI agent that automatically designs aptamers by assembling the target-binding fragments in the pool, and named it AiDTA (Ai-driven docking-then-assembling). In the design process, AiDTA used the MCTS algorithm to explore the fragment assembly action spaces by generating simulated aptamer sequences, and then trained a policy-value neural network that predicts the assembly probabilities of the fragments and the values of the corresponding assembled aptamers, thereby guiding the agent to generate aptamers with similar secondary structures of the constituent fragments. To verify AiDTA, we designed DNA aptamers against four important disease-related protein targets: the receptor-binding domain (RBD) of the spike protein of SARS-CoV-2 Omicron B.1.1.529 variant^35^, the spike RBD of SARS-CoV-2 Omicron BA.2.86 variant^36^, TNF-like ligand 1A (TL1A)^37^, and the A1 domain of von Willebrand factor (vWF-A1)^38^. For each target, 6∼9 designed aptamers were synthesized and then validated for their binding activities by MicroScale Thermophoresis (MST) measurements. The MST results showed that most of the designed aptamers exhibited target-binding affinities with equilibrium dissociation constants (*K*d) in the nanomolar range. These results strongly suggest that we have successfully achieved the de novo design of protein-binding aptamers for the first time.

## Results

### 1. Library of short nucleic acid fragments

In our DTA study^24^, the target OA was a marine biotoxin with a relatively small size, so we used only four mononucleotides (i.e., A, C, G, and T) as the short nucleic acid fragments for molecular docking. As a result, the manually designed aptamers had a relatively simple secondary structural element (SSE). However, previous studies have shown that during the SELEX screening, the used nucleic acid libraries that contain certain SSEs often yielded better results, suggesting that the SSEs are important components of aptamers^39,40^. Because proteins are significantly larger in size than OA, the structures of aptamer-protein complexes clearly demonstrated that the SSEs in aptamers frequently participate in their binding to the target proteins. For example, such a binding pattern can be found in the complex structures from the Protein Data Bank (PDB) codes: 3HXO^41^, 4PDB^42^, and 6SY6^43^.

Based on the above consideration, besides the short single-stranded fragments, we also constructed secondary structural fragments for the molecular docking. Specifically, we used the 4 single nucleotides (A, G, C, and T) to construct the single-stranded fragments with lengths of 3, 4, 5, and 6 nucleotides, and the double-stranded fragments with base-pair lengths of 3, 4, 5, and 6 base pairs (Fig. 1b). To avoid intensive computation by too many fragments, the lengths of the constructed fragments were not greater than 6 nucleotides or 6 base pairs. Consequently, we constructed a library containing 5,440 short single-stranded and 5,440 double-stranded fragments. We then predicted and optimized their 3D structures using the 3dRNA/DNA program and their sequences and corresponding secondary structural states. To ensure that the fragment structures are in the lowest-energy conformations, we further minimized their energies using the AMBER99bsc1 force field in AmberTools23 (Methods)^44,45^. After energy minimization, we obtained a library of short nucleic acid fragments for the molecular docking against a given target protein. Selected examples of the 3D structures of the single-stranded and double-stranded fragments are shown in Figs. 1c and d, respectively.

### 2. Fragment docking based on protein target structure

At the initial phase of this study, Omicron B1.1.529 variant was the prevalent SARS-CoV-2 strains. Thus, we selected the receptor-binding domain (RBD) of its spike protein as the first target protein of this study. Based on the structures of RBD in complex with antibodies (PDB codes: 7QTK^35^ and 7QNW^46^), we defined its antibody-binding interface as those amino acids within 6 Å of any atom of the antibodies (Fig. 2a) and considered the interface as the desired binding epitope of designed aptamers^47^.To determine the high-affinity binding sites for each fragment on the target protein, we performed global molecular docking for all the fragments in the library against RBD using HDOCK^48^. Examples of the docked fragments are shown in Fig. 2b. We then calculated the distances between the geometric centers of the highest-scoring conformations of each docked fragment and the Cα atoms of amino acids in the desired epitopes, and those fragments with a distance < 6 Å were defined as epitope-binding fragments. As a result, approximately 7,000 epitope-binding fragments were obtained, as illustrated in Fig. 2c.

**Fig. 2:**
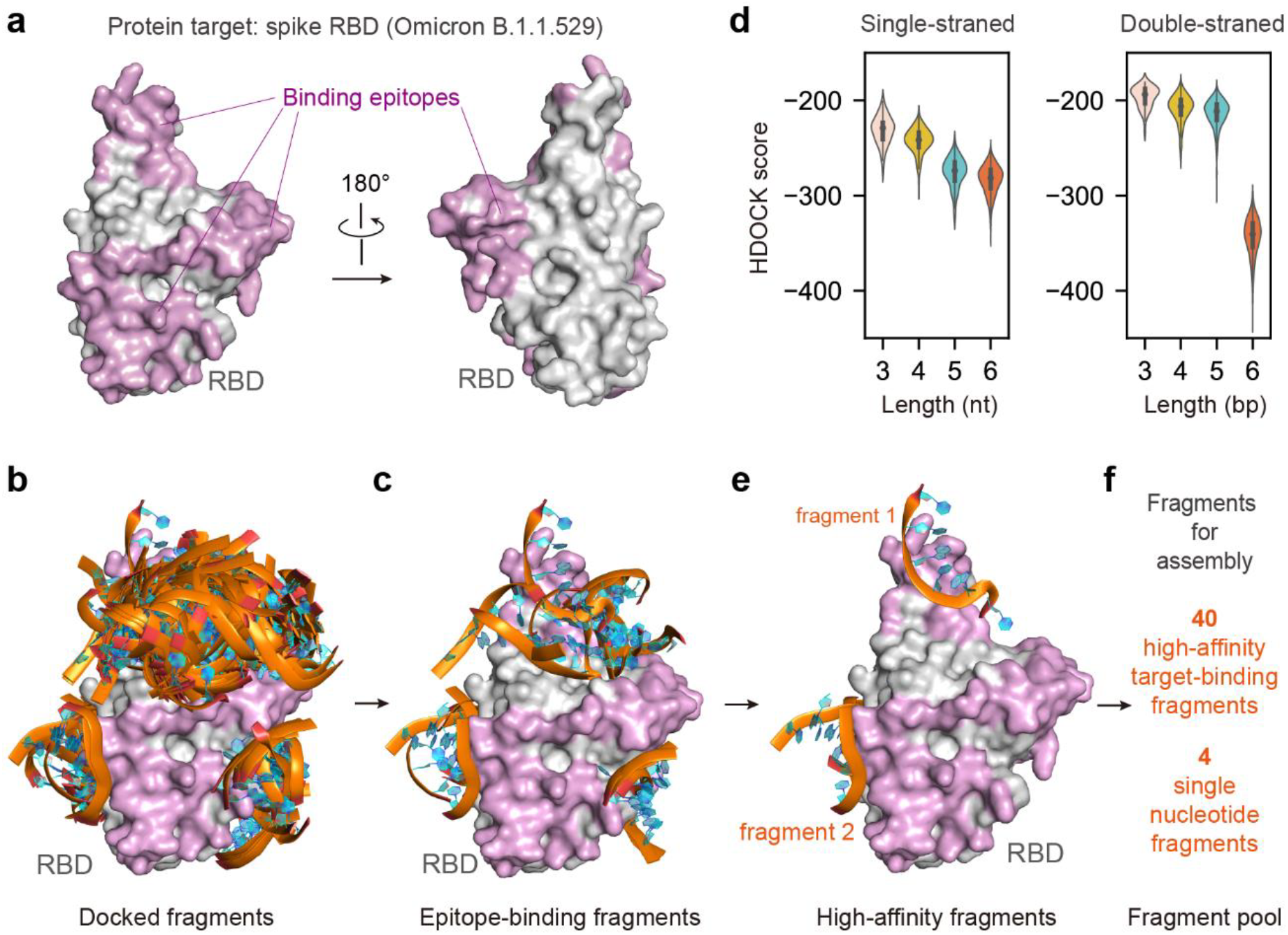
Protein target and molecular docking of short nucleic acid fragments. **a**, Structure of the SARS-CoV-2 spike RBD (Omicron B.1.1.529) and binding epitope identified based on the antibody-RBD interfaces in the complex structures (PDB codes: 7QTK, 7QNW). **b**, Binding modes of different fragments to RBD obtained by fragment docking using HDOCK. Phosphate groups of nucleotides are shown in orange, and pentose sugars and bases are shown in cyan. **c**, Epitope-bound fragments. **d**, Docking score distributions of the single-stranded and double-stranded fragments of different lengths. **e**, Selected examples of high-affinity fragments filtered by binding sites and docking scores. **f**, The fragment pool contained 4 single-nucleotide fragments and 40 high-affinity target-binding fragments.

Next, we analyzed the HDOCK docking scores of those epitope-binding fragments, as shown in Fig. 2d. The docking scores of the single-stranded fragments ranged from -200 to - 350, and those of the double-stranded fragments ranged from -150 to -400. To ensure high-affinity binding, we ranked the fragments by the docking scores in the ascending order and then selected the top 5 single-stranded or double-stranded fragments of each length (nt or bp), resulting in 40 fragments. These fragments were regarded as potential binding fragments with high affinity (Fig. 2e). The docking scores and their secondary structures of these fragments are listed in Tables S1 and S2. According to the binding confidence defined by HDOCK, the binding confidence of the filtered single-stranded fragments ranged from 93% to 98%, and that of the double-stranded fragments ranged from 94% to 99%. Therefore, these 40 fragments formed a pool of fragments for the subsequent aptamer assembly. Similar to our previous study^24^, we also incorporated the four mononucleotides (A, C, G, and T) into the fragment pool to ensure that any two discrete target-binding fragments could be connected by a certain number of single nucleotides of various lengths in the assembly process. Thus, the assembly fragment pool contained 44 fragments (Fig. 2f).

### 3. Fragment assembly by the AI agent AiDTA

To select suitable fragments from the fragment pool and then assemble them into a full-length aptamer capable of binding to the target, we developed an AI agent by combining RL, the MCTS algorithm, and the deep neural networks, as indicated by the workflow in Fig. S1. As shown in Fig. 3a, AiDTA starts with the initial null sequence. Subsequently, at steps *t* = 1, 2, …, *T*, AiDTA iteratively selects a fragment from the pool to append or insert into the assembled sequence at the last step, finally forming a continuous aptamer sequence of *l* nucleotides. In the process, the same fragments can be selected repeatedly in different steps. Therefore, the assembly action at step *t, a*_t_, consists of two continuous operations: 1) selection of a fragment among the *N*_frag_ available fragments; 2) then, selection of a linkage site among the *N*_site_ possible positions; consequently, the number of assembly actions is *N*_frag_ × *N*_site_ (Fig. 3b). To describe the RL environment state of AiDTA, i.e., the assembled sequence, we used a matrix *M* with dimensions 6 × *L* to encode the sequence (Fig. S2). Columns represent the nucleotide sequence positions, and *L* is the maximum length of the designed aptamers. Rows 1-6 represent the four nucleotides at the given position, and their base-pairing states in the original constituent fragments.

**Fig. 3:**
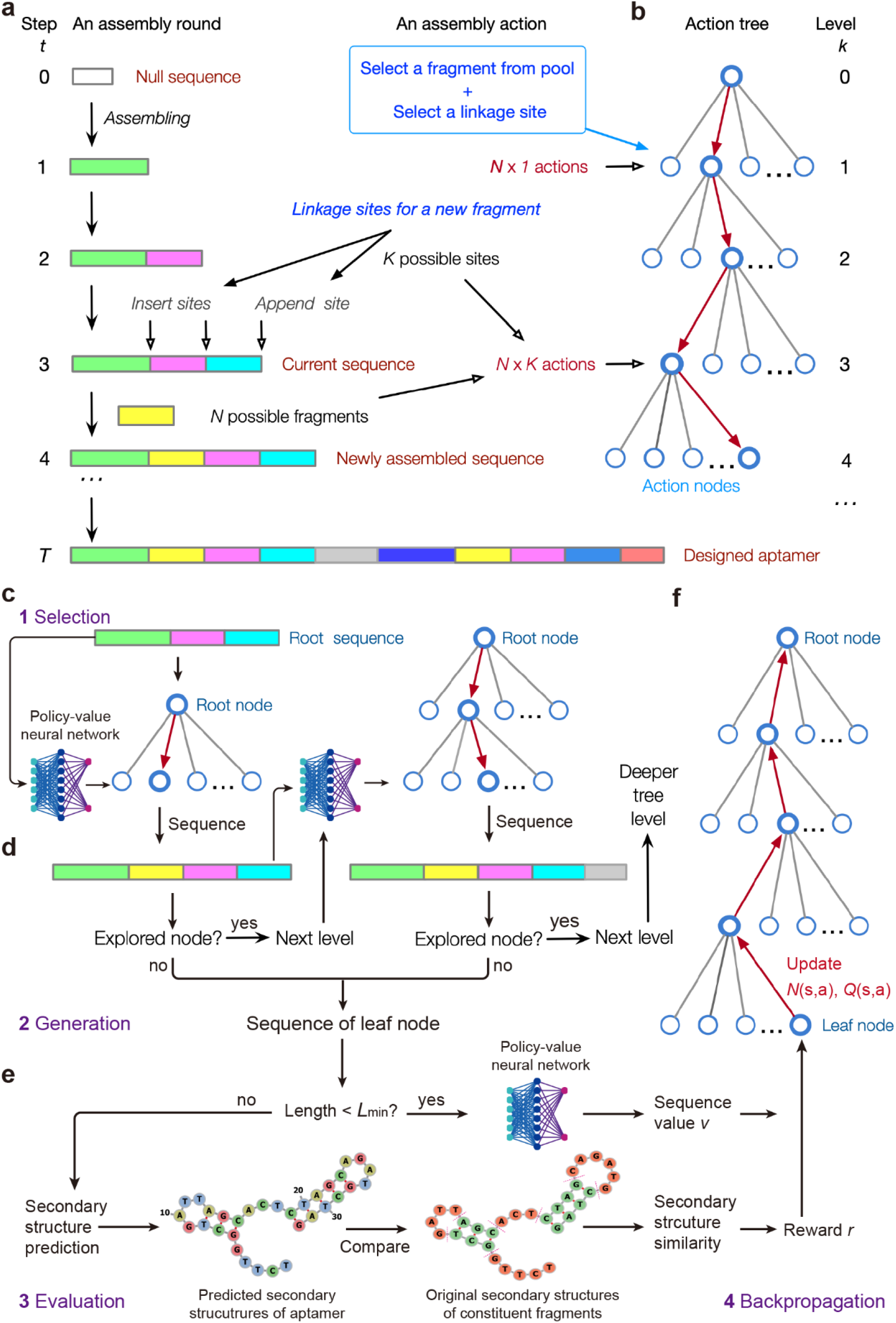
De novo aptamer design using AiDTA. **a**, Fragment assembly process. At each assembly step *t*, a fragment is selected from the 44-fragment pool and either inserted or appended at a specific site (i.e., linkage site) until the assembly round is completed. At each step, *N* fragments and *K* linkage sites are considered. **b**, Assembly action tree. The fragment assembly decision process starts at the root node (level *k* = 0) and branches into possible assembly actions at deeper levels (*k* = 1, 2, …). **c**, Action node selection. AiDTA navigates the action tree by evaluating both explored and unexplored nodes using the policy-value neural network. **d**, Sequence generation. When reaching an unexplored node (leaf node) during simulations, AiDTA generates the corresponding leaf sequence. **e**, Sequence evaluation. Based on whether the sequence length exceeds the minimum threshold *L*_min_, AiDTA either uses the policy-value neural network to predict the sequence value, or determines the reward by secondary structure similarity calculation, in which the predicted secondary structures of an aptamer are compared with those of the original constituent fragments indicated by the slash lines in red, and the reward is determined by the similarity value. **f**, Backpropagation update. Node values and visit counts are updated along the backpropagation tree path from the leaf node to the root node.

Similar to AlphaZero^31^, AiDTA uses the encoding matrix of a given sequence *s* as the input to the policy-value neural network: *f*_θ_(*s*). This deep neural network consists of a convolutional module, residual modules, and an output module (Fig. S3). Its policy head outputs the prior selection probabilities of all possible assembly actions from *s*: ***p***(*s*); and its value head outputs the value of the sequence: *v*. During the assembly process, AiDTA uses the MCTS algorithm to generate simulation sequences for training the neural network, in turn, the trained network guides the MCTS algorithm to efficiently explore optimal fragment assembly actions. At step *t*, AiDTA treats the assembled sequence *s*_t-1_ from the last step as the current root sequence, and considers it as the root node of the assembly action tree at level *k* = 0 (Fig. 3b); then, AiDTA performs 100 independent MCTS assembly simulations (Figs. 3c-f, Fig. S4). Each MCTS simulation consists of four sub-steps (phases):

1. *Selection of assembly action*: Starting with the current *s*_root =_ *s*_t-1_, AiDTA generates simulation sequences by exploring the action tree to deeper levels (*k* = 1, 2, 3, …). The nodes of the action tree correspond to different assembled sequences of different lengths derived from *s*_root_. For a given node defined by the sequence *s*, to determine its target node for moving, AiDTA uses the policy-value neural network to predict the selection probabilities of actions ***p***(*s*_k_), and then calculates the cumulative average value of action *Q*(*s*_k_,*a*_k_) and the exploration bonus *U*(*s*_k_,*a*_k_) (Methods). AiDTA selects the action node with the maximum value as the target node, 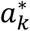 (Fig. 3c).
2. *Generation of sequence*: If the sequence defined by the target node has been explored in any previous simulation, AiDTA continued to explore the action tree towards nodes at deeper levels according to sub-step 1. If not, AiDTA expanded the tree to the target node as a leaf node and the generated its sequence *s*_leaf_ (Fig. 3d).
3. *Evaluation of sequence*: If the length of *s*_leaf_ was less than the desired minimum length *L*_min_ (50 nt in this study), its value was evaluated directly using the policy-value neural network with *s*_leaf_ as the input. Otherwise, AiDTA used RNAstructure^49^ to predict the secondary structure of *s*_leaf_, and calculated the secondary structure similarity between the aptamer sequence and the original constituent fragments (Methods). Higher similarity implies that the secondary structures of the aptamer resemble those of the constituent fragments, and thereby the assembled aptamer likely has similar protein-binding modes. An aptamer sequence with a similarity value ≥ 0.9 was considered as a successful assembly and got a reward value of +1; otherwise, the reward value is -1 (Fig. 3e).
4. *Backpropagation of action tree*: After evaluating *s*_leaf_, AiDTA traced back from the leaf node to the root node. Each ancestor node along the tree path increments its visit count *N*(s,a) and recursively updates *Q*(s,a) according to the value or reward of *s*_leaf_ (Fig. 3f).

After 100 simulations, AiDTA calculated the probability of each assembly action under *s*_root =_ *s*_t-1_. The action with the maximum sum of probability distribution and Dirichlet noise was then selected to generate the actual assembled sequence *s*_t_ (Fig.S4). The introduction of Dirichlet noise could ensure a balance between exploration and exploitation of action combinations throughout the assembly process.

### 4. Sequence analysis of the assembled aptamers

As demonstrated in Fig. S5, after about 500 epochs of training, AiDTA assembled about 20,000 aptamer sequences with a value of secondary structure similarity ≥ 0.9. To investigate their secondary structure features, we first analyzed the similarities of their secondary structures with the original constituent fragments. For comparison, we also generated 20,000 sequences of 50-61 nucleotides in length by randomly assembling the fragments in the pool. As shown in the similarity distributions in Fig. 4a, the secondary structure similarities of the AiDTA aptamers are higher than those of the random sequences. Significantly certain numbers of aptamers generated by AiDTA have a similarity value ranging from 0.9 to 1.0, but those of almost all the random sequences were less than 0.9. This demonstrates the good performance of AiDTA in assembling sequences with similar structures of the original constituent fragments.

**Fig. 4:**
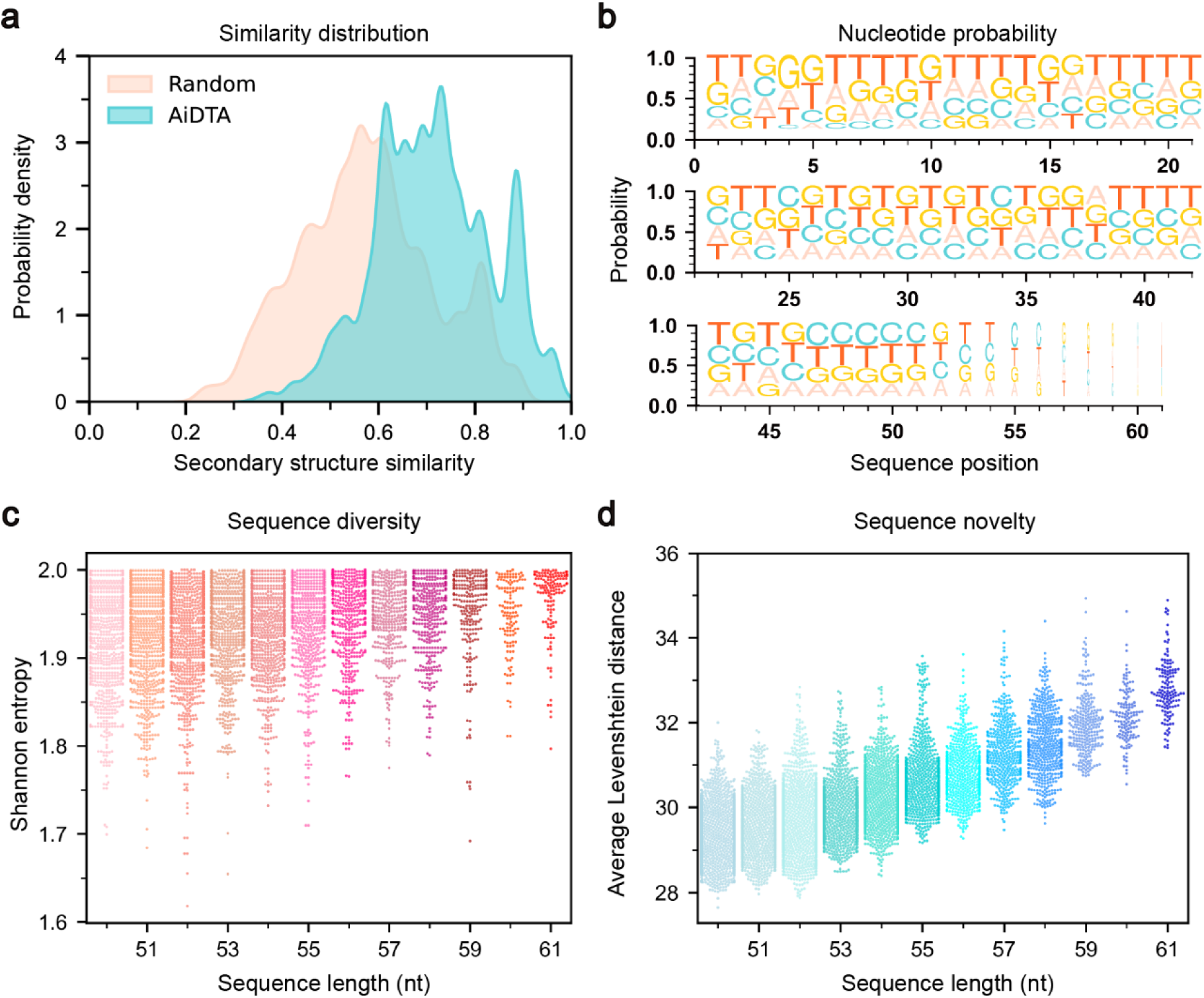
Sequence evaluation of AiDTA-designed aptamers. **a**, The Ai-DTA designed aptamers showed a 15% higher secondary structure similarity compared to the randomly assembled sequences. **b**, The nucleotide compositions (A, C, G, T) in the designed aptamers. **c**, Shannon entropies of the designed aptamers of various lengths. **d**, Average Levenshtein distances of the designed aptamers of various lengths.

Since sequence diversity reflects the ability of AiDTA to explore the fragment assembly space, we further analyzed the diversity and novelty of the assembled sequences. We visualized these sequences using WebLogo and found that for the AiDTA aptamers of 50-61 nt, the frequencies of the four nucleotides at each position were nearly equal (Fig. 4b). To quantify their diversity, we calculated the Shannon entropies of the nucleotides for each sequence. As shown in Fig. 4c, the average Shannon entropy of the assembled sequences reached 1.8, close to the theoretical maximum of 2.0, indicating that the probabilities of the four nucleotides occurring were approximately equal. In addition, we also examined the novelty of the sequences by calculating their Levenshtein edit distances. As illustrated in Fig. 4d, almost all sequence pairs had an average Levenshtein edit distance greater than 28, indicating significant sequence differences between them. Thus, AiDTA is able not only to assemble aptamer sequences that preserve the original secondary structures of the constituent fragments, but also to ensure high diversity and uniqueness among these sequences. In other words, AiDTA can explore a large fragment assembly space and generate aptamers with low sequence homology.

### 5. Site-specific filtering of aptamers for experimental verification

To enrich the aptamers capable of binding to the target protein, we conducted a multi-step filtering. First, we predicted the 3D structures of the assembled aptamers using 3dRNA/DNA and performed energy minimization to optimize their conformations, as the examples shown in Fig. S6. Based on the optimized structures, we then calculated the Euclidean distance and surface geodesic distance between the 5’ and 3’ ends of each aptamer. Similarly, we determined the maximum Euclidean and geodesic distances for the amino acid residues in the target epitopes (Figs. 5a and b; see also Methods). If both the maximum Euclidean and geodesic distances of a given aptamer are greater than those of the protein epitopes, this aptamer was considered to cover the epitopes. Using this criterion, approximately 15,000 aptamers were filtered as candidates. Next, we used HDOCK to perform global docking for these aptamers against the protein. For each aptamer, the docked complex structure with the highest docking score was selected as the representative conformation of the given aptamer in complex with the protein (Fig. S7). Then, the Euclidean distances between the aptamer nucleotides and the epitope residues were calculated; if anyone of the nucleotide-residue distances was less than 7 Å, this aptamer was considered to potentially bind to the epitope (Fig. 5c). Approximately 8,000 aptamers were selected in this way.

**Fig. 5:**
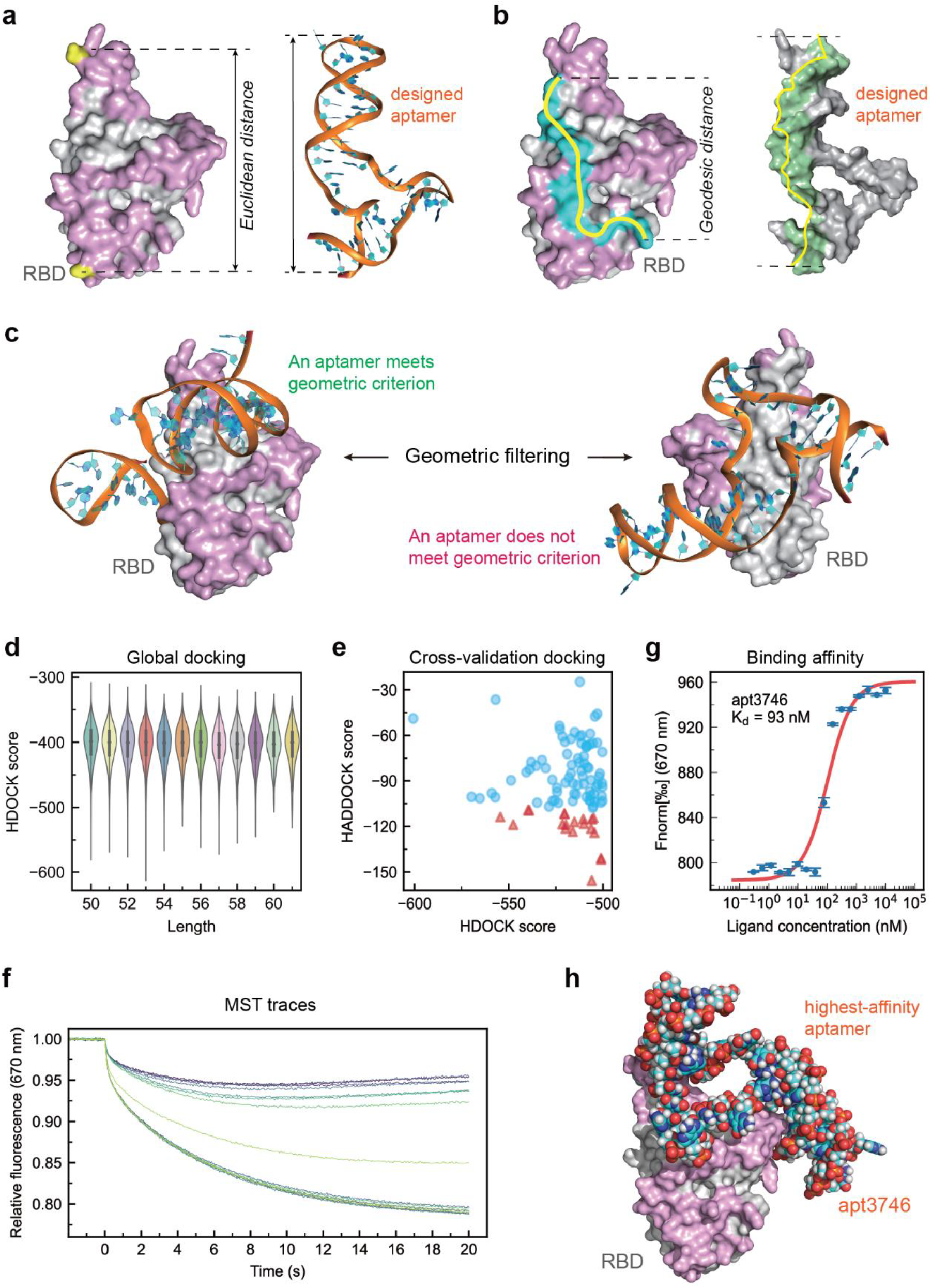
Geometric filtering of designed aptamers and experimental validation. **a**, Maximum Euclidean distance between two epitope residues and that between two nucleotides of a designed aptamer. **b**, Corresponding geodesic distance between two epitope residues and that between two nucleotides of a designed aptamer. **c**, Geometric filtering of designed aptamers based on the Euclidean and geodesic distances. **d**, Distributions of HDOCK docking scores of designed aptamers. **e**, Cross-validation HADDOCK docking scores of designed aptamers. **f**, MST measurements of aptamer-target protein interactions with varying aptamer concentrations and a fixed protein concentration. **g**, Equilibrium dissociation constant (*K*_d_) of the highest-affinity aptamer measured by MST. **h**, Predicted 3D structure of the highest-affinity aptamer in complex with protein target.

Subsequently, the binding capabilities of these aptamers were further evaluated by cross-validation docking. To this end, the representative complexes of the top 100 aptamers ranked by the HDOCK docking scores were validated by global docking using HADDOCK^50^ (Figs. 5d, e). The top 20 aptamers ranked by the HADDOCK docking scores were then identified. To further narrow the aptamers for experimental verification, we used AlphaFold3 (AF3)^51^ to predict the complex structures for these 20 aptamers based on their sequences and the protein. The complex structures predicted by AF3 confirmed that 6 aptamers were bound to the desired epitope of the target protein. Thus, these six aptamers were synthesized for the subsequent experimental verification of their binding activities.

### 6. Experimental verifications by MST

To verify the binding activity of a given aptamer, we dissolved the synthetic aptamer in dry powder form in 1 mL of 1 × PBS solution, and then measured its concentration in solution by using the NanoDrop spectrophotometer. Based on the concentration, we further diluted it to 100 nM and aliquoted it into 16 Eppendorf tubes. To ensure a uniform folding of the aptamers in solution, we used the Eppendorf thermocyclers to heat the aptamer sample to 95°C and maintain it for 5 minutes, then performed an annealing process by cooling at a rate of 1°C per minute until it reached 25°C. Similarly, we measured the concentration of the protein solution and diluted it to 20,000 nM, and then performed 15 rounds of 2-fold serial dilutions, resulting in 16 solution samples of different concentrations. Next, equal volumes of the aptamer solutions were added to each protein solution sample, respectively. After incubation at 25°C with slow shaking for 1 hour, the mixed solutions were aspirated using a MO-K022 capillary for the MST measurements.

To determine the binding affinities, we loaded the sample capillaries into the Monolith X instrument and measured the changes in fluorescence intensity resulting from aptamer-protein interactions, as illustrated by an example in Fig. 5f. Each MST measurement for an aptamer and the target protein was performed in triplicate, and the equilibrium dissociation constant (*K*_d_) was calculated using the MO.Control 2 software based on the average fluorescence shift values from the three replicates. As expected, four out of the six tested aptamers exhibited *K*_d_ values below 1 μM. Notably, two aptamers displayed the very high affinity, with *K*_d_ values of 93.0 and 94.4 nM, respectively (Fig. 5g, Fig. S7). For comparison, we also determined the *K*_d_ value of the SELEX-derived aptamer CoV2-RBD-1 by MST, which was 173 nM (Fig. S7). The structure of the highest-affinity aptamer apt3746 in complex with the target protein is shown in Fig. 5h, while the complex structures of the other tested aptamers are presented in Fig. S8. These MST results demonstrate that AiDTA successfully designed nucleic acid aptamers capable of binding to the target protein with high affinity.

### 7. Aptamers targeting other protein targets

To test the application potential of AiDTA, we further designed aptamers for other three important disease-related target proteins: the spike RBD of SARS-CoV-2 Omicron BA.2.86 variant, TNF-like protein 1A (TL1A), and the A1 domain of von Willebrand factor (vWF-A1) (Figs. 6a-c).

**Fig. 6:**
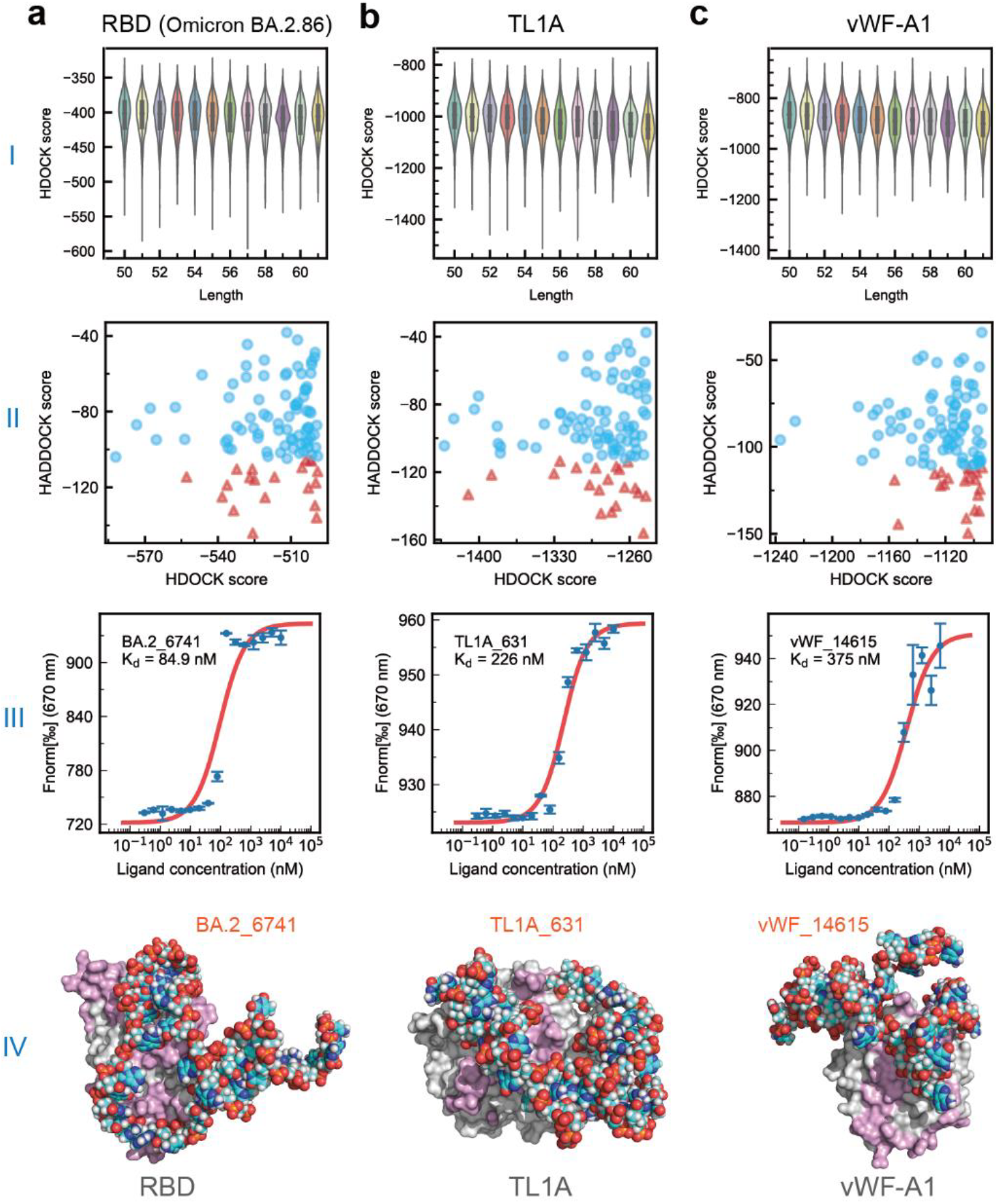
Experimental validations of aptamers designed for other protein targets. **a**, Results for the SARS-CoV-2 spike RBD (Omicron BA.2.86). (I) HDOCK docking scores of designed aptamers; (II) Cross-validation HADDOCK docking scores for the top-ranked aptamers using HDOCK; (III) MST-measured equilibrium dissociation constant (*K*_d_) of the highest-affinity aptamer; (IV) Predicted 3D model of the highest-affinity aptamer in complex with protein target. **b**, Results for TL1A (I-IV corresponding to panels in **a**). **c**, Results for vWF-A1 (I-IV corresponding to panels in **a**).

As mentioned earlier, this study initially focused on the spike RBD of the Omicron B.1.1.529 variant. However, with the continuous evolution of SARS-CoV-2, the Omicron BA.2.86 variant emerged as the new dominant strain. Yang et al. demonstrated that this variant exhibits significant antigenic changes and potential immune escape capabilities^52^. Additionally, Zhang et al. found that BA.2.86 can efficiently infect cells and evade neutralizing^53^, highlighting the urgent need to develop detection and therapeutic strategies against rapidly evolving variants^54^. Based on the structure of the antibody-spike complex (PDB code: 8QTD^55^), we identified the target epitope on the RBD for aptamer design and utilized AiDTA to design eight aptamers for MST validation. The MST results revealed that six aptamers exhibited *K*_d_ values below 1 μM, with three of them showing *K*_d_ values < 100 nM, including the highest-affinity aptamer (BA.2_6741) with a *K*_d_ of 84.9 nM (Fig. 6a and Fig. S9). These high-affinity aptamers demonstrate that AiDTA can rapidly design aptamers targeting viral variants without the need for experimental screening, providing a powerful tool for the rapid discovery of protein-binding aptamers with potential applications in antiviral detection and therapeutic development.

TL1A plays a critical role in inflammatory diseases such as inflammatory bowel disease (IBD) and eosinophilic asthma through its interaction with the receptor DR3^56^. IBD includes Crohn’s disease and ulcerative colitis (UC). A recently completed Phase II clinical trial demonstrated that the monoclonal antibody Tulisokibart significantly improved the clinical symptoms in UC patients when used to treat UC^57^. Another study showed that the TL1A/DR3 signaling pathway significantly enhances the activation and effector functions of ILC2s, leading to eosinophil recruitment and exacerbated airway inflammation^37^. These findings highlight the importance of developing drugs to modulate the TL1A/DR3 signaling pathway or methods to detect the TL1A protein. To design binding aptamers targeting TL1A, we used the TL1A-DCR3 complex structure (PDB code: 3K51^58^) and that of TL1A-DR3 complex predicted by AF3 to identify the binding epitope on TL1A and then designed eight aptamers. MST experiments showed that four aptamers had *K*_d_ values < 1 μM, and two of them had *K*_d_ values < 300 nM (Fig. S10). The best aptamer (TL1A_631) had a *K*_d_ value of 226 nM (Fig. 6b).

Finally, we employed AiDTA to design aptamers targeting vWF-A1, a potential diagnostic marker and therapeutic target for thrombotic diseases, inflammatory vascular diseases, tumor metastasis^59,60^. The development of vWF-A1-binding aptamers could enable precise modulation of its activity to address diverse disease states, serve as robust tools for assessing vascular disease risk and progression, and provide a foundation for more accurate and effective diagnostic and therapeutic strategies for vascular diseases. Indeed, several RNA and DNA aptamers targeting vWF-A1 have been discovered through SELEX, some of which have advanced to clinical trials^61^. Here, we tested the capability of AiDTA by designing new aptamers for vWF-A1. Based on the SELEX-derived aptamer ARC1172 and its complex structures with vWF-A1 (PDB codes: 3HXO^41^ and 7F49^62^), we defined the target epitope on vWF-A1 and designed seven aptamers for experimental validation. The MST results revealed that three aptamers exhibited *K*_d_ values below 1 μM, with the highest-affinity aptamer (vWF_14615) displaying a *K*_d_ value of 375 nM (Fig. 6c). For comparison, the *K*_d_ value of the SELEX-derived aptamer ARC1172, as determined by MST, was 453 nM (Fig. S11).

## Discussion

In this study, we used the DeepRL strategy to develop an AI agent for *de novo* design of nucleic acid aptamers targeting proteins, named AiDTA. In our AI-driven design approach, we first docked the short nucleic acid fragments to the protein to identify the high-affinity target-binding fragments, and then used AiDTA to automatically assemble these fragments into the full-length aptamers. To validate AiDTA, we designed DNA aptamers for four important protein targets and experimentally verified their binding affinities using MST. For each target protein, we synthesized fewer than 10 designed aptamers for experimental validation. The MST results demonstrated that most of the designed aptamers exhibited target-binding affinities with *K*_d_ values in the nanomolar range, and the highest-affinity aptamers achieved *K*_d_ values below 100 nM. When compared to traditional multi-round SELEX screening, our AiDTA approach is significantly cost-effective and precise.

Since Ellington and Szostak first introduced the SELEX method in 1990^15^, significant advancements have been made in aptamer discovery. To address the limitations of the original method, various modified SELEX approaches have been developed, such as cell-SELEX^63^, microfluidic chip-SELEX^64^, and hydrogel-SELEX^65^, etc. However, even compared to the improved SELEX methods, AiDTA still represents a transformative leap in nucleic acid aptamer discovery. AiDTA uses DeepRL to automate aptamer design, eliminating the multi-round selection steps inherent in SELEX. This dramatically improves efficiency: whereas traditional SELEX requires multiple experimental isolation and purification steps, AiDTA only requires the synthesis and validation of a small number of test samples, significantly reducing cost and process complexity. More importantly, the AiDTA design process is highly precise and can be focused on predefined binding sites for the target protein. This site-specific design approach avoids the waste of resources associated with non-site-specific binding sequences in the SELEX process.

Unlike previous *in silico* aptamer design methods, AiDTA designed aptamers directly from computationally identified target-binding fragments without relying on any experimental screening data. For instance, Raptgen^21^ and AptaDiff^22^ require SELEX data particular for a given target in order to train and optimize their AI models. Consequently, the trained models are, in principle, mainly suitable for the given target. Recently, RhoDesign designed RNA aptamers that target small molecules without SELEX data; however, this approach still requires predetermined aptamer structures related to the target. In contrast, AiDTA does not require any predetermined aptamer structures, making it a more general approach to the de novo design of protein-binding aptamers. Indeed, AiDTA has been validated for multiple protein targets, including virus-related proteins, human inflammatory factors, and coagulation factors, demonstrating its applicability for various protein targets. In the four demonstrated targets, two of them already had SELEX aptamers. Compared to the tested samples in their SELEX screening, our approach only synthesized a few of tested aptamers for validation, fully demonstrating the value of our approach to discovery of protein-binding aptamers. Thus, our method is extremely helpful for designing affinity aptamers for large numbers of protein targets, such as in aptamer-based precision proteomics^66^. To the best of our knowledge, AiDTA is the first computational *de novo* design method for protein-binding aptamers.

As a proof of concept, this study primarily focused on developing AiDTA. Further research is certainly needed to improve the method in the future. Since our approach is the first of its kind, currently no similar *in silico* design methods that can be used for systematic benchmarking against it. Also, in our study the protein targets were selected simply for the convenient demonstration of the AiDTA design process, rather than for the potential applications of the designed aptamers. Thus, our study lacked functional characterization of the developed aptamers. More tests focusing on actual applications are needed for additional protein targets. Moreover, AiDTA may be improved by integrating advanced structural prediction models, such as AlphaFold3, to design aptamers more directly based on the 3D structures of protein targets. In addition, combining AiDTA with high-throughput experimental validation platform may automate the entire process from computational design to experimental validation. Furthermore, exploring the potential of AiDTA in designing aptamers against small molecules and in developing multivalent aptamers could create new opportunities for its application in various fields.

In summary, we have developed a DeepRL-based agent AiDTA that achieved the de novo design of protein-binding aptamers for the first time. Experimental validation demonstrated that AiDTA-designed aptamers targeting disease-related proteins exhibit high binding affinities. Remarkably, AiDTA achieves a very high success rate compared to the labor-intensive experimental screening. As an efficient and cost-effective alternative to the experimental screening, AiDTA establishes a completely new, general framework for the rational design of aptamers. Our study opens a new avenue for obtaining protein-binding aptamers, thereby advancing potential aptamer applications in biopharmaceutical development and molecular diagnostics.

## Methods

### Prediction of nucleic acid structures

The RNAstructure program^49^ was used to predict the secondary structure of single-stranded DNA sequences that are output as CT format files. The CT files were then converted to Dot-Bracket Notation (DBN) files using the ct2dot conversion tool. The 3dRNA/DNA program^67^ was used to predict the 3D structures of the aptamers based on their sequences and the secondary structures defined the DBN files. The predicted structures were then optimized by its built-in Monte Carlo simulated annealing module of 3dRNA/DNA.

### 3D structure refinements

The 3D structures predicted by 3dRNA/DNA were further refined through energy minimization using the ParmBSC1 force field in AmberTools23^44,45^. During the minimization, the implicit solvent model igb5 was used as the solvent. The cutoff distance for short-range electrostatic and van der Waals interactions was set to 14 Å. The maximum number of minimization cycles is set to 5000, and the steepest descent algorithm was used for the first 500 steps, followed by a switch to the conjugate gradient algorithm.

### Encoding of sequences and representation of assembly actions

The assembled aptamer (or environmental state) of AiDTA was encoded by a matrix with dimensions 6 × *L* (Fig. S2). Columns represent the nucleotide sequence positions of the aptamer, with *L* being the maximum length. Row 1 to 4 represent the four nucleotides at the given position, and rows 5 and 6 denote their base-pairing states in the constituent fragments. In the matrix, A, G, C, and T in the sequence are represented as 1, 2, 3, and 4, respectively; brackets “(“and “)” are represented as 5, and a period “.” is represented as 6.

The assembly action space at step *t* is represented as a one-dimensional vector of *N*_frag_ × 50, where 50 is the maximum number of linkage sites in *s*_t_. The assembled aptamer (or state) *s*_t_ is updated iteratively until its sequence length exceeds the desired minimum length, *L*_min_. In the assembly process, corresponding reward of the assembled sequence *s*_t_ (0 < *t* ≤ *T*) was defined as *r*_t_ (0 < *t* ≤ *T*).

### Architecture of policy-value neural network

The policy-value neural network consists of a convolutional module, seven residual modules, a policy module, and a value module (Fig. S3). The initial convolutional module comprises a 3×3 convolution kernel with 6 input channels and 256 output channels, using a stride of 1 and padding of 1. After the convolution operation, the output is sequentially processed by batch normalization (BatchNorm) and a ReLU activation function. The network utilizes seven residual blocks, each consisting of the output from the previous module and two convolutional sub-modules. Each convolutional sub-module has 256 input and output channels, employs a 3×3 kernel, and uses a stride of 1 and padding of 1. The output of each convolutional operation undergoes batch normalization followed by a ReLU activation.

The policy module includes a 1×1 convolutional kernel with 256 input channels and 16 output channels, using a stride of 1. After the convolution, the output is processed by batch normalization and a ReLU activation function. Subsequently, a fully connected layer transforms the output into a vector of 2200 dimensions. The convolutional kernel of the value module has 256 input channels and 8 output channels, using a 1×1 kernel and a stride of 1. The output is processed by batch normalization and ReLU activation, followed by a fully connected layer that converts the output into a 256-dimensional vector, which is then passed through another ReLU activation. Finally, this vector is reduced to a scalar through another fully connected layer, and the output is mapped to the range [-1, 1] using the Tanh function.

### MCTS simulation and neural network training

One MCTS simulation corresponds to iterative assembly steps from the root sequence node *s*_root_ to a leaf node *s*_leaf_ in the action tree, where each node represents a specific assembled sequence *s* derived from *s*_root_. As shown in Figs. 3c-f, the simulation process include the following phases^31,68^:

1. *Selection*: Starting from *s*_root_, AiDTA explored child nodes at the deeper levels until it reaches an unexpanded node (unexplored level), which represents a sequence that has not yet been assembled in the simulations. At each node, the next assembling action *a*^∗^ for the given sequence *s* is determined by the PUCT (Polynomial Upper Confidence Trees) formula^30^:

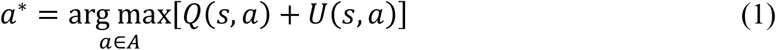

where *A* is the action set, *Q*(*s, a*) is cumulative average value of action *a*, and *U*(*s, a*) is exploration bonus:

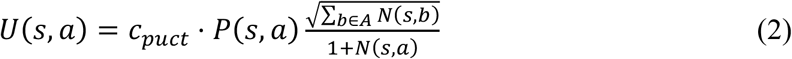

*c*_*puct*_ is a constant that balances exploration and exploitation, *P*(*s, a*) is the prior probability of action *a* predicted by the policy-value neural network, and *N*(*s, a*) is the current visit count for the action *a* from *s, ∑*_*b*∈*A*_ *N*(*s, b*) is the total visit count for all possible actions from *s*.
2. *Expansion*: When an unexpanded node is encountered, the selected action *a* is executed on the current sequence *s* to a new fragment assembling, resulting in a child sequence *s*^′^. This newly assembled sequence *s*^′^ is then added to the action tree as the leaf node of this simulation.

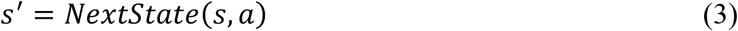
3. *Evaluation*: For the leaf sequence *s*^′^with a sequence length < *L*_min_, its value *v* and action probabilities ***p***(*s*^′^) are calculated by the policy-value neural network, *f*_*θ*_:

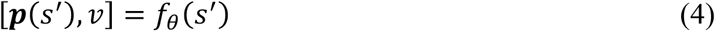

For a sequence with a length ≥ *L*_min_, its reward *r* is calculated as in the following subsection.
4. *Backpropagation*: After a simulation, the visit count and cumulative average value from *s*_leaf_ to *s*_root_ are updated as:

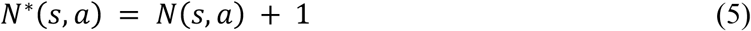

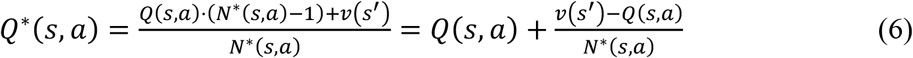

where *N*^∗^(*s, a*) is the updated count after an additional visit, and *Q*^∗^(*s, a*) is the updated *Q*(*s, a*) with the value of the new sequence *s*^′^.

After the simulations from the root sequence *s*_root_, the actual action at time step *t* is sampled based on the policy **π**_t_, which is proportional to the 1/τ power of the visit counts at step *t*:

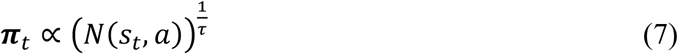

At the end of each assembly round, AiDTA generates (*s*_*t*_, **π**_t_, r_t_) for each sequence *s*_t_ encountered, recording the actual choices **π**_t_ and the rewards *r*_t_. AiDTA stores these data in a buffer to train the policy-value neural network according to the following loss function:

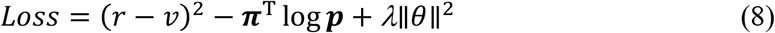

where *r* is the reward of the assembled sequence, and the (*r* − *v*)^2^ term measures the difference between the predicted value *v* and the actual reward *r* from the simulated aptamers. The ***π***^T^ log ***p*** term is the cross-entropy loss of the policy function that quantifies the consistency between the MCTS probabilities ***π***^T^ and the actual probabilities ***p***. *θ* is the model weights, and *λ* is the parameter of L2 weight regularization.

### Secondary structure similarity and reward of an assembled aptamer

The secondary structure of an assembled aptamer was predicted using RNAStructure^49^. AiDTA then compares the secondary structure state of each nucleotide to the original state of the corresponding nucleotide in the given constituent fragment. If the secondary structure sates are identical, the similarity score for the given nucleotide is 1; if they differ, the score is 0. The secondary structure similarity for the assembly aptamer is defined as the sum of the nucleotide scores divided by the sequence length, i.e., the average similarity score per nucleotide. The higher the similarity, the more likely it is that the assembled aptamer retains the high-affinity binding modes of the constituent fragments. An assembled aptamer with a similarity value ≥ 0.9 receives a reward *r* = 1, while those with a similarity value below 0.9 receive a reward *r* = -1.

### Sequence analysis of assembled aptamers

For each assembled aptamer, the probability *p*_*i*_ of each of the four nucleotides is calculated. The Shannon entropy of each sequence is then used to characterize the diversity of nucleotide types within the aptamer:

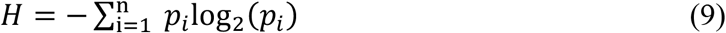

Levenshtein distance is used to measure the difference between two aptamers (strings). Specifically, it represents the minimum number of edit operations required to transform one aptamer sequence (string) into the other, including insertion, deletion, and substitution of characters.

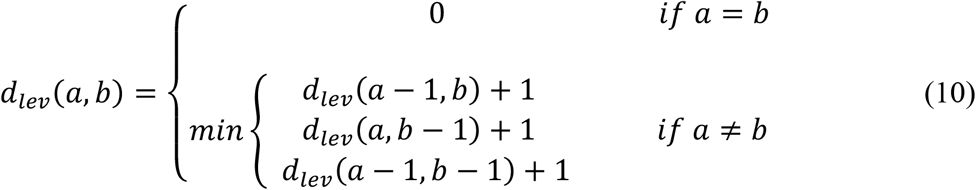

Furthermore, the average Levenshtein distance for a collection of *n* strings is the arithmetic mean of all pairwise Levenshtein distances. If *s*_1_, *s*_2_, …, *s*_*n*_ denote the strings (aptamer sequences) in the set, the average Levenshtein distance for a given string (i.e., aptamer) *s*_*i*_:

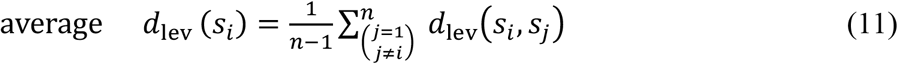

### Molecular docking for aptamer-protein pairs

The HDOCK program^48^ (version 1.1) was used to predict the complex structure of an aptamer with its protein target using the *ab initio* free docking protocol. The PDB file of the protein was set as the receptor, and the PDB file of a given aptamer was set as the ligand. To predict the binding complex, HDOCK performed global docking for the default number of times and outputted the complex files in the .out format, which can be converted into PDB file by the creapl tool. The built-in ITScorePP function of HDOCK was used as the docking score to evaluate nucleic acid-protein interactions. The binding confidence score defined in HDOCK served as an indicator of the likelihood that the given aptamer and protein would bind: a confidence score > 0.7 indicates a high probability of binding, a score between 0.5 and 0.7 suggests potential binding, and a score < 0.5 indicates unlikely binding.

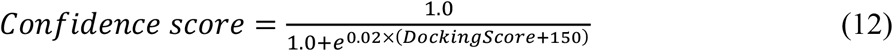

The HADDOCK program^50^ was used to perform cross-validation docking of the aptamer-protein complexes. FreeSASA was first employed to calculate the solvent-accessible surface area (SASA) of each amino acid and nucleotide, identifying those with a SASA > 40 Å^2^. These residues were then input into the *active-passive-to-ambig*.*py* script of HADDOCK to generate the ambiguous restraints file for the given protein and aptamer. In the docking configuration file, the protonation states of protein histidine residues were automatically defined, and a dielectric constant of 78 was used. During docking, random removal of residues was disabled, and ambiguous constraints were enabled, along with additional features such as random ambiguous interaction constraints, center-of-mass constraints, and surface constraints. Finally, 1,000 docking sampling runs were performed for each aptamer-protein pair, and the default HADDOCK score was used to rank the docking poses, with the weight of desolvation energy set to 0, as recommended for protein-nucleic acid docking.

### Aptamer synthesis and MST measurements

The DNA aptamers were synthesized by Sangon Biotech (Shanghai) Co., Ltd. After synthesis, the Cy5 fluorescent dye was conjugated to the 5’ end of the aptamers through a chemical coupling reaction. Following the coupling reaction, the aptamers were purified, and the labeling efficiency was verified to ensure successful fluorescent labeling. Subsequently, the aptamers were further purified using high-performance liquid chromatography (HPLC), and the target aptamers were precisely collected via a UV detector, followed by lyophilization to ensure high purity and long-term stability, providing high-quality aptamer samples for subsequent MST experiments.

MST experiments were conducted using the Monolith x system to quantify the binding affinity between an aptamer and its target protein. The IR settings were automatically configured, with each sample measurement consisting of a 5-second interval followed by a 20-second laser-on time. The Cy5-labeled aptamer was dissolved in PBS buffer (8.1 mmol/L Na_2_HPO_4_, 1.5 mmol/L KH_2_PO_4_, 137 mmol/L NaCl, 2.7 mmol/L KCl) and heated at 95°C for 10 minutes in a PCR machine. The temperature was then gradually reduced at a rate of 1°C per minute until reaching 25°C. The target protein was serially diluted in a twofold gradient, starting from a maximum concentration of 20,000 nM down to 0.61 nM, resulting in 16 distinct concentrations. Equal volumes of the aptamer and target protein solutions were mixed and incubated on a shaking platform at 180 rpm for 1 hour. The mixtures at the 16 concentrations were then loaded into capillaries (Cat# MO-K022) for MST measurements. Independent experiments were performed at least three times to determine the average equilibrium dissociation constant (*K*_d_) for the aptamer-protein interaction.

## Data availability

All datasets used for the model training and evaluation are publicly available, with the processing details described in Methods.

## Code availability

The source codes for AiDTA are publicly available via GitHub at https://github.com/Fudan-HQLab/AiDTA.

## Acknowledgements

This work was supported by the National Key Research and Development Program of China (2021YFA0910604), the National Natural Science Foundation of China (31971377, 31671386).

## Author contributions

Q.H. supervised the project. G.G. and Q.H. conceived the project. G.G. developed the computer programs, and G.G., J.Q., X.Q. and L.W. analyzed the computer codes. G.G and L.G. conducted the computational design, and G.G., L.G. and X.H. carried out the experimental validation. G.G. and Q.H. drafted and revised the manuscript. All authors contributed to the drafting and revision of the manuscript.

## Additional information

**Supplementary information** is available for this paper at …

**Correspondence and requests for materials** should be addressed to Q.H.

