## Supplementary Information for "De novo design of protein-binding aptamers through deep reinforcement learning assembly of nucleic acid fragments"

**Table S1 | Top five single-stranded fragments for binding to the Spike RBD of SARS-CoV-2 Omicron B.1.1.529 ranked by the HDOCK docking scores**

| Length (nt) | Fragment ID | Sequence | Secondary structure | HDOCK score |
|---|---|---|---|---|
| 3 | 3_6 | AGG | ... | -273.33 |
| | 3_17 | GAA | ... | -256.64 |
| | 3_19 | GAC | ... | -253.80 |
| | 3_22 | GGG | ... | -259.97 |
| | 3_46 | CTG | ... | -263.03 |
| 4 | 4_64 | ATTT | .... | -276.40 |
| | 4_69 | GAGA | .... | -295.26 |
| | 4_105 | GCCA | .... | -285.71 |
| | 4_126 | GTTG | .... | -274.32 |
| | 4_238 | TCTG | .... | -275.69 |
| 5 | 5_965 | TTAGA | ..... | -327.28 |
| | 5_968 | TTAGT | ..... | -327.53 |
| | 5_981 | TTGGA | ..... | -325.20 |
| | 5_984 | TTGGT | ..... | -326.29 |
| | 5_1015 | TTTGC | ..... | -324.94 |
| 6 | 6_3231 | TACGTC | ...... | -345.67 |
| | 6_3838 | TCTTTG | ...... | -347.52 |
| | 6_4022 | TTCTGG | ...... | -345.65 |
| | 6_4081 | TTTTAA | ...... | -347.54 |
| | 6_4083 | TTTTAC | ...... | -346.81 |

**Table S2 | Top five double-stranded fragments for binding to the Spike RBD of SARS-CoV-2 Omicron B.1.1.529 ranked by the HDOCK docking scores**

| Length (bp) | Fragment ID | Sequence | Secondary structure | HDOCK score |
|---|---|---|---|---|
| 3 | d_3_2 | AAG&CTT | (((&))) | -212.66 |
| | d_3_23 | GGC&GCC | (((&))) | -232.89 |
| | d_3_35 | CAC&GTG | (((&))) | -214.36 |
| | d_3_40 | CGT&ACG | (((&))) | -221.74 |
| | d_3_52 | TAT&ATA | (((&))) | -221.74 |
| 4 | d_4_13 | AATA&TATT | ((((&)))) | -256.19 |
| | d_4_58 | ATCG&CGAT | ((((&)))) | -258.33 |
| | d_4_118 | GTGG&CCAC | ((((&)))) | -249.75 |
| | d_4_193 | TAAA&TTTA | ((((&)))) | -251.85 |
| | d_4_201 | TACA&TGTA | ((((&)))) | -262.16 |
| 5 | d_5_94 | AGGTG&CACCT | (((((&))))) | -297.49 |
| | d_5_350 | GGGTG&CACCC | (((((&))))) | -299.57 |
| | d_5_606 | CGGTG&CACCG | (((((&))))) | -298.02 |
| | d_5_647 | CCAGC&GCTGG | (((((&))))) | -298.92 |
| | d_5_902 | TCAGG&CCTGA | (((((&))))) | -290.35 |
| 6 | d_6_144 | AACATT&AATGTT | ((((((&)))))) | -431.29 |
| | d_6_1168 | GACATT&AATGTC | ((((((&)))))) | -427.58 |
| | d_6_1369 | GGGGCA&TGCCCC | ((((((&)))))) | -426.86 |
| | d_6_2905 | CTGGCA&TGCCAG | ((((((&)))))) | -425.24 |
| | d_6_3310 | TATCTG&CAGATA | ((((((&)))))) | -436.13 |

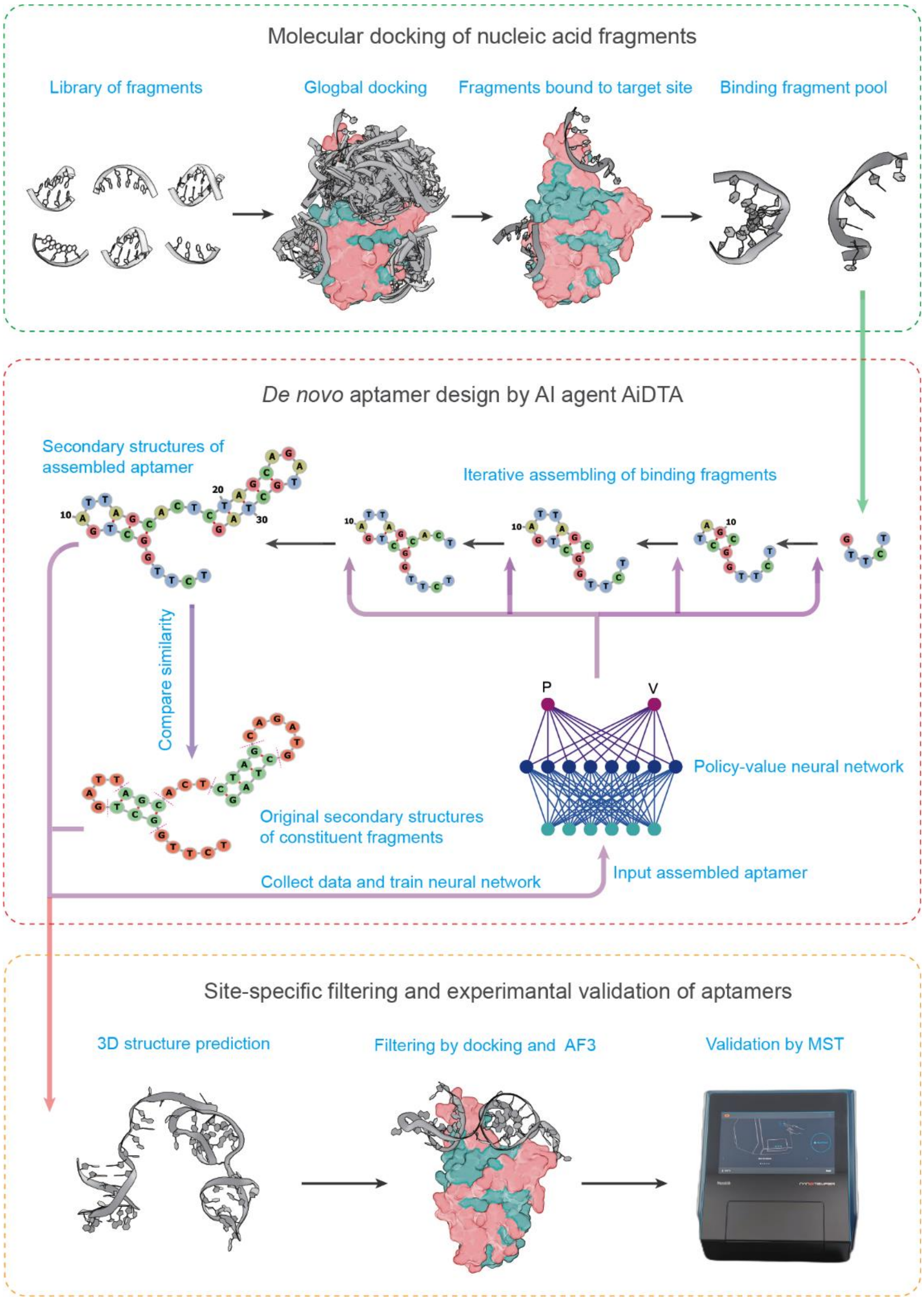


**Fig. S1 | Workflow of *de novo* aptamer design using the AI agent AiDTA.** The process comprises three key steps: (1) construction of a fragment library containing short single-stranded and double-stranded nucleic acid fragments, (2) AiDTA-driven assembly of fragments into aptamers, and (3) site-specific filtering of designed aptamers and experimental validations.

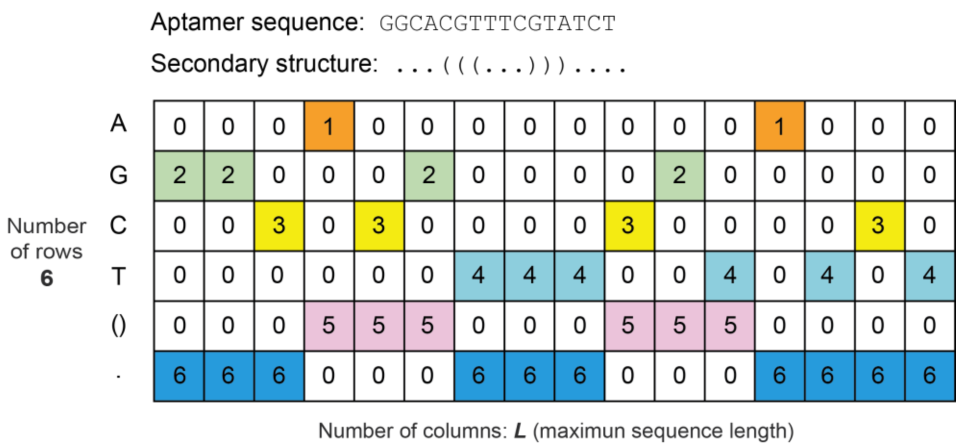


**Fig. S2 | Encoding of an aptamer sequence using a matrix.** The matrix *M* has dimensions of 6 × L, where L is the maximum length of aptamer sequences. The six rows are divided into two parts: rows 1–4 represent the four nucleotides (A, C, G, T), and rows 5–6 indicate the single-stranded or double-stranded state of the nucleotides.

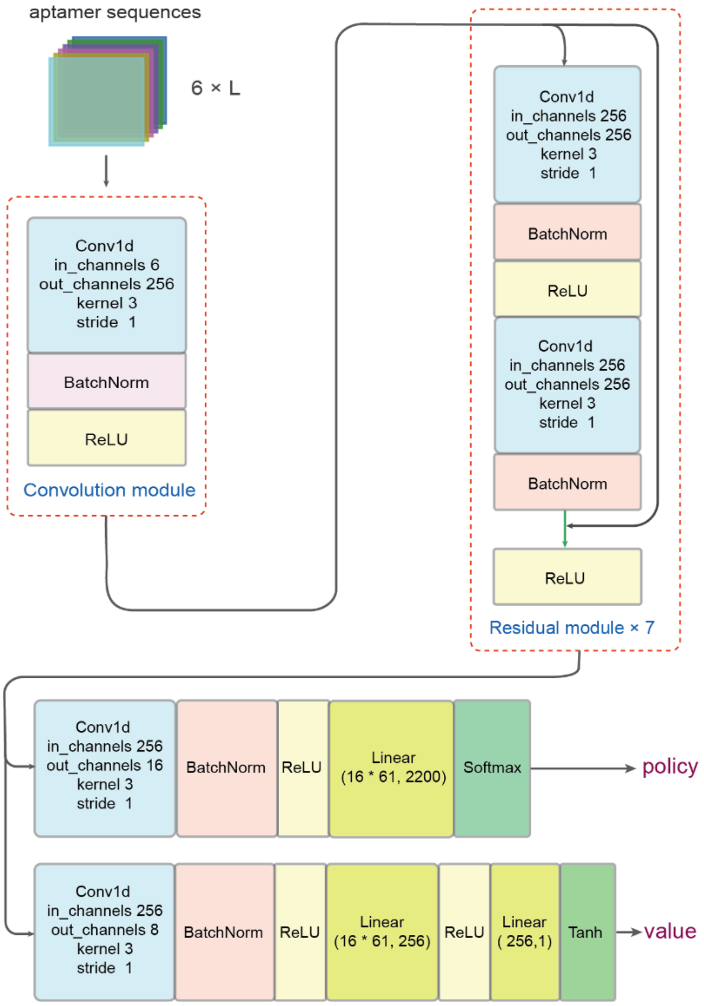


**Fig. S3 | Architecture of policy-value neural network.** The deep neural network consists of four components: a convolutional model, a residual module, a policy head, and a value head.

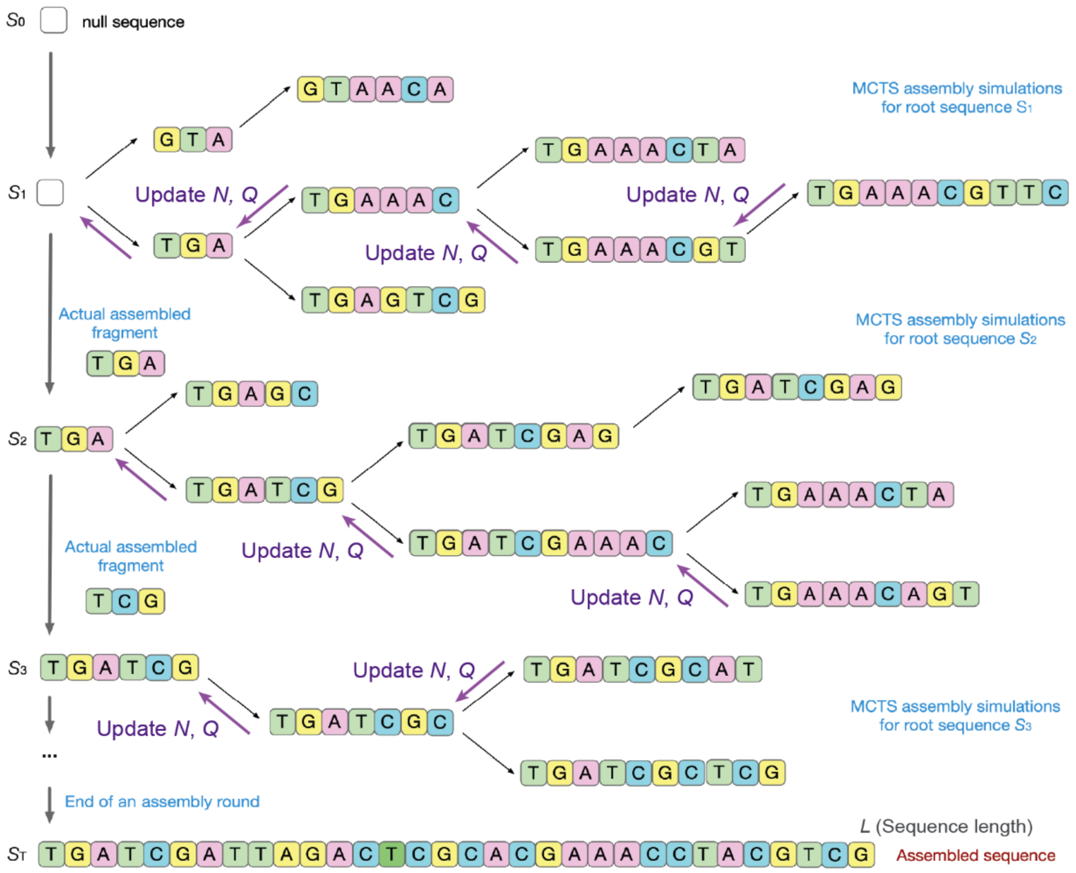


**Fig. S4 | MCTS simulations of fragment assembly in AiDTA.** At each assembly step $t$ ($t = 1, 2, ..., T$), AiDTA first performs 100 fragment assembly simulations via the MCTS algorithm, exploring multiple action pathways and testing different fragment combinations. Each MCTS simulation initiates from the corresponding root sequence at step $t$ and terminates at an unexplored leaf node. After completing the simulations at step $t$, AiDTA determines the actual fragment for assembly and updates both the node visit counts ($N$) and $Q$-value. The assembly process continues until the generated aptamer sequence exceeds the minimum length threshold $L_{\mathrm{min}}$.

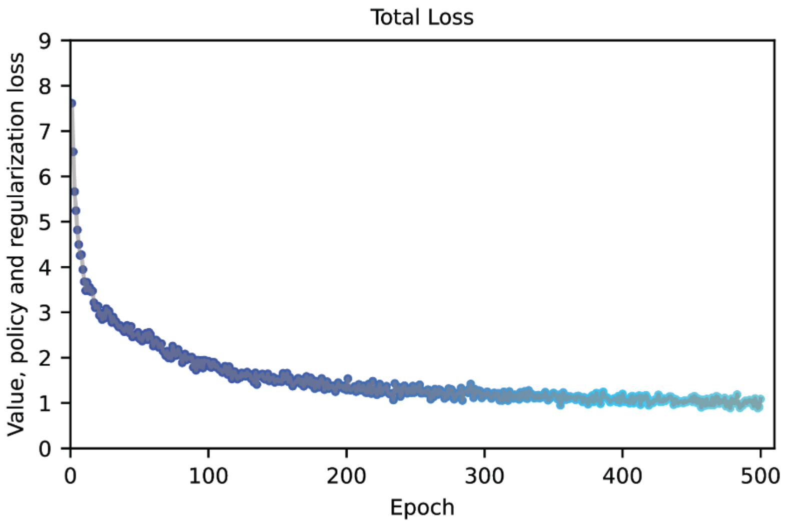


**Fig. S5 | Total loss of the policy-value neural network during AiDTA training**. As the training iterations progress, the total loss (comprising value loss, policy loss, and regularization loss) exhibits a consistent and progressive decline, indicating improved performance and convergence of the neural network toward optimal parameters.

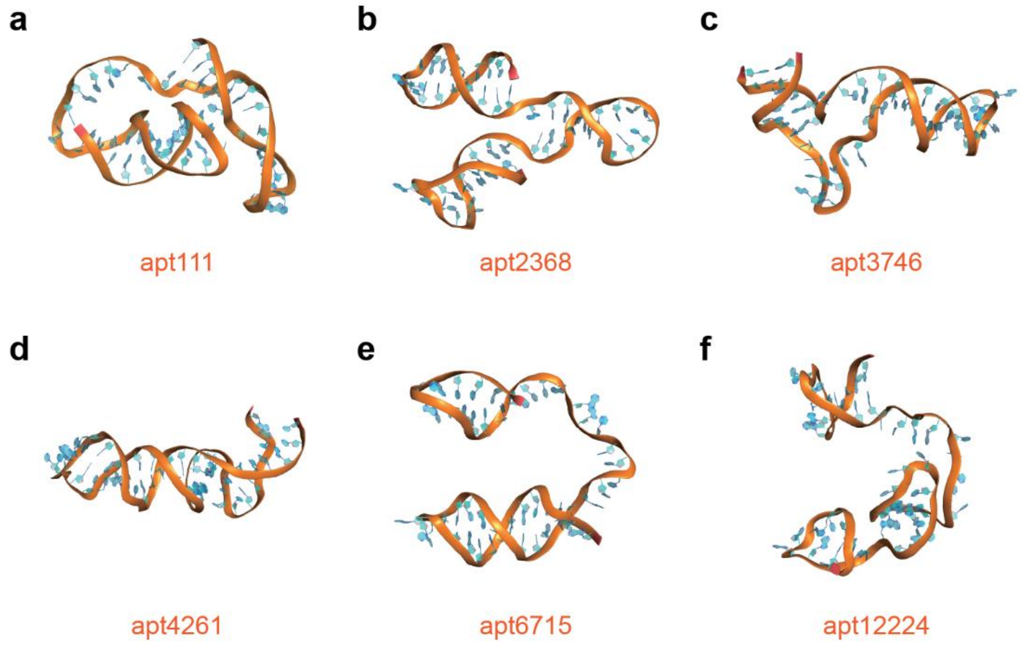


**Fig. S6 | Predicted 3D structures of designed aptamers using 3dDNA/RNA**. **a,** apt111**. b,** apt2368**. c,** apt3746**. d,** apt4261. **e,** apt6715**. f,** apt12224**.**

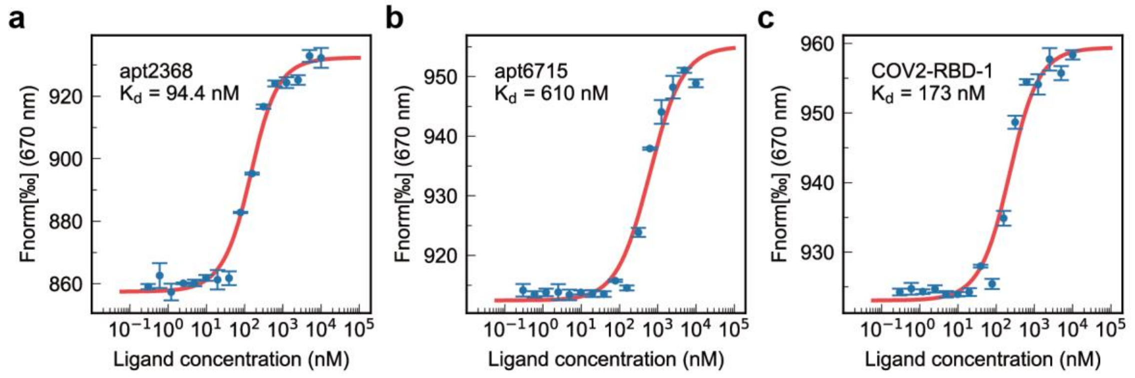


**Fig. S7 | MST results of other designed aptamers with nanomolar binding affinities targeting the Spike RBD of SARS-CoV-2 Omicron B.1.1.529.** **a**, apt2368. **b**, apt6715. **c**, CoV2-RBD-1. Note that CoV2-RBD-1 was obtained using SELEX by Song et al., *Anal. Chem*. **2020**, 92:9895–9900.

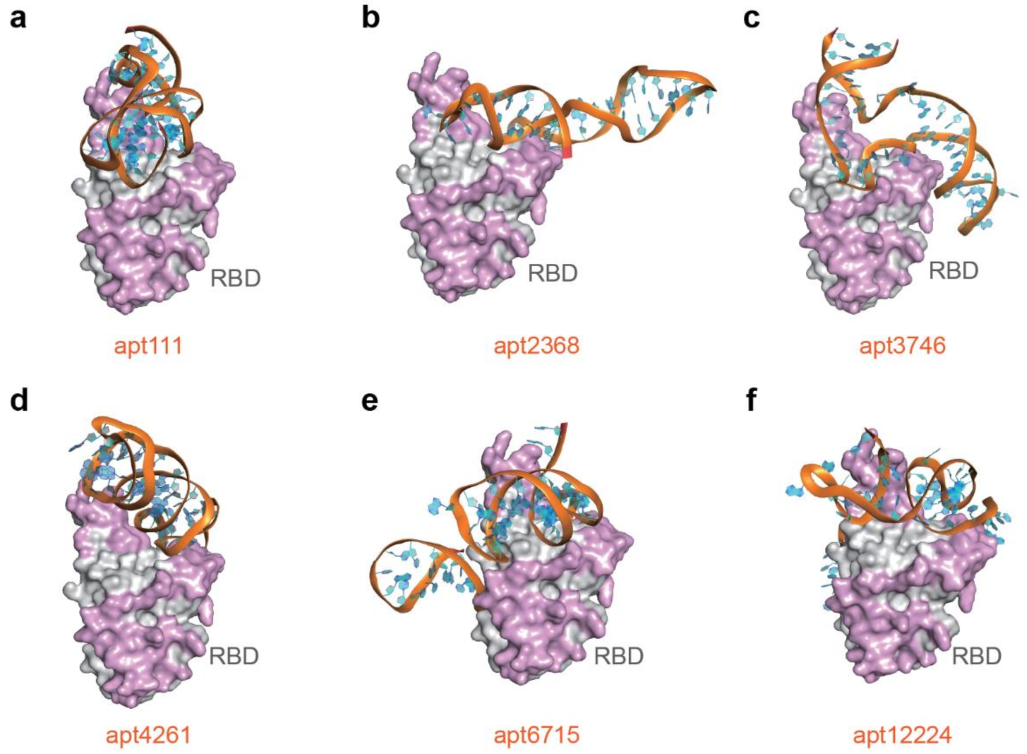


**Fig. S8 | Predicted structures of designed aptamers in complex with the Spike RBD of SARS-CoV-2 Omicron B.1.1.529 using HDOCK.** **a**, apt111. **b**, apt2368. **c**, apt3746. **d**, apt4261. **e**, apt6715. **f**, apt12224.

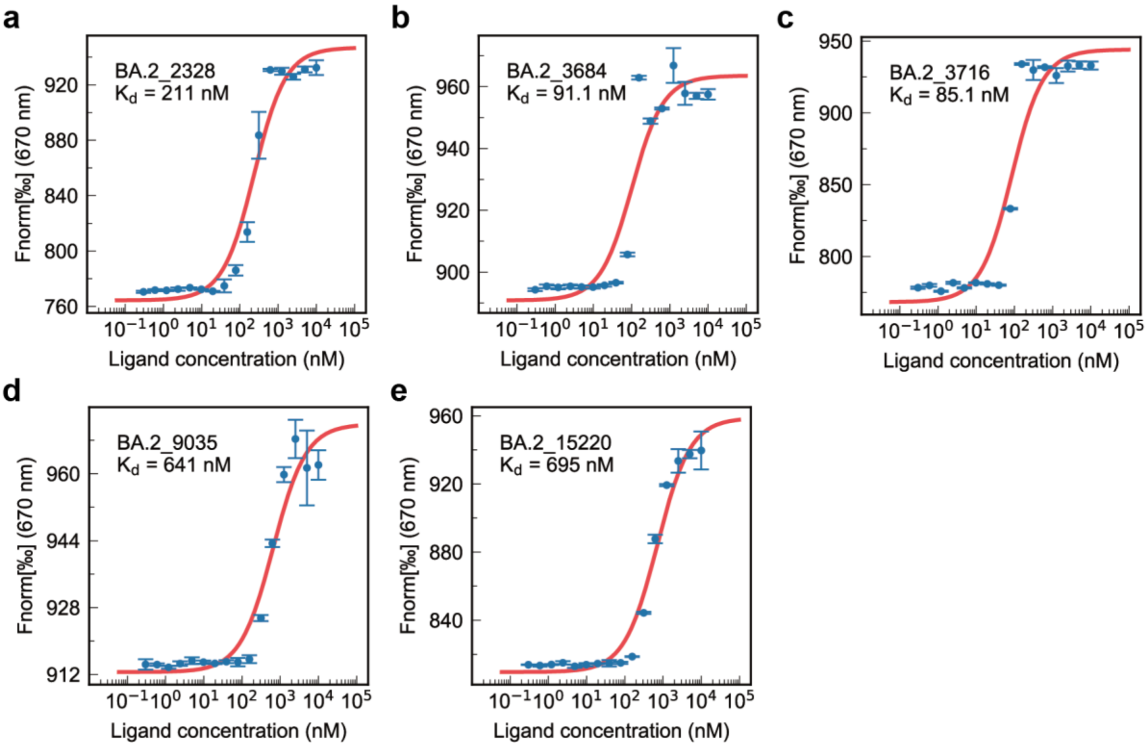


**Fig. S9 | MST results of other designed aptamers with nanomolar binding affinities targeting the Spike RBD of SARS-CoV-2 Omicron BA.2.86. a**, BA.2_2328 **b**, BA.2_3684. **c**, BA.2_3716. **d**, BA.2_9035. **e**, BA.2_15220.

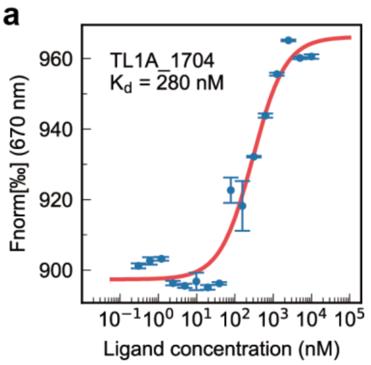


**Fig. S10 | MST results of other designed aptamers with nanomolar binding affinities targeting TL1A. a**, TL1A_1704.

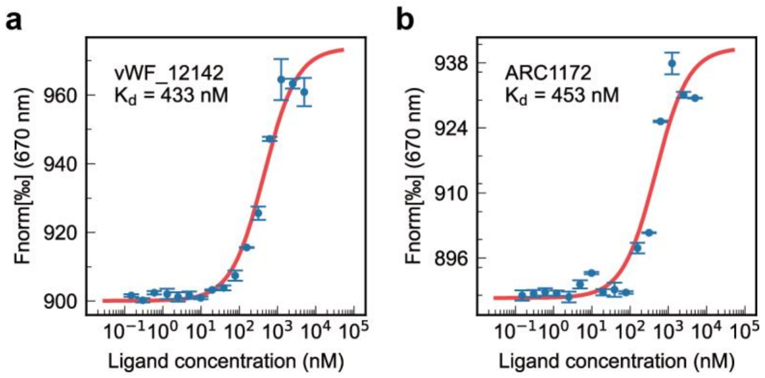


**Fig. S11 | MST results of other designed aptamers with nanomolar binding affinities targeting vWF-A1. a**, vWF_12142. **b**, ARC1172. Note that ARC1172 was obtained using SELEX by Huang et al., *Structure* **2009**, 17*:*1476–1484.
